# Amino Acid Insertion Energetics in a POPC Bilayer from Unbiased Molecular Dynamics

**DOI:** 10.64898/2026.05.07.723583

**Authors:** Sophie Bories, Patrick Lague

## Abstract

Membrane association is governed by the thermodynamics of amino acid partitioning between water and the lipid bilayer. Here, we quantified amino acid side-chain insertion energetics in a 1-palmitoyl-2-oleoyl-*sn*-glycero-3-phosphocholine (POPC) bilayer using unbiased molecular dynamics simulations. Equilibrium depth distributions of 28 analogs, including multiple protonation states, were converted into potentials of mean force (PMFs) by Boltzmann inversion. The resulting PMFs reproduced the main features of bilayer partitioning. Hydrophobic analogs favored the bilayer core, aromatic analogs were stabilized in interfacial regions, and polar or charged analogs remained unfavorable in the hydrophobic interior. A diglycine analog representing the peptide backbone behaved similarly to uncharged polar residues. Depth-dependent pK_a_ profiles and orientational analyses further showed how protonation equilibria and aromatic-ring alignment influence insertion energetics. Agreement with experimental hydrophobicity scales supports the robustness of the approach. These results provide an efficient and internally consistent framework for characterizing bilayer insertion energetics and establish a reference for future studies in more complex lipid environments.

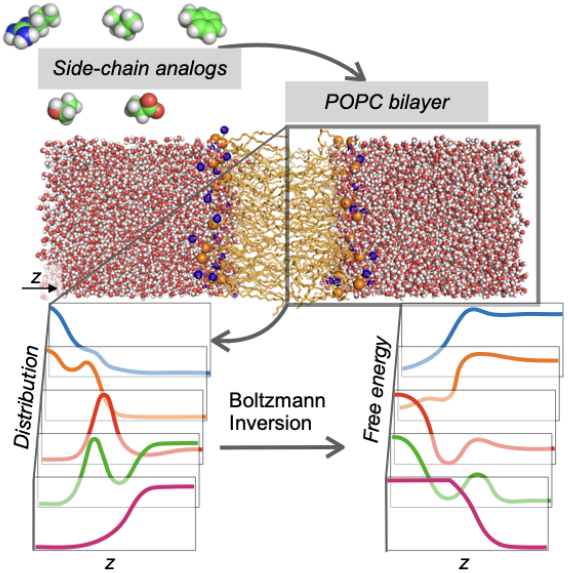

**SIGNIFICANCE:** Membrane-associated proteins represent a large fraction of the proteome and include many major drug targets, yet quantitative understanding of their interactions with lipid bilayers remains limited. Here, we present an unbiased molecular dynamics framework for systematically determining amino acid side-chain insertion free energies in a model bilayer. By deriving potentials of mean force directly from equilibrium depth distributions, this approach enables internally consistent comparisons across residue classes and protonation states without biasing restraints. The resulting free-energy profiles reproduce established hydrophobicity trends and show how protonation equilibria and aromatic-ring orientation modulate bilayer partitioning. This scalable strategy provides a quantitative reference for residue-level membrane thermodynamics and establishes a foundation for extending insertion energetics to more diverse lipid compositions and more complex membrane-associated systems.

## INTRODUCTION

Membrane proteins and membrane-associated proteins constitute a substantial fraction of the human proteome and account for more than half of approved drug targets (1–3). Their function depends critically on interactions with biological membranes, whose lipid composition and physicochemical properties vary across organisms and cellular compartments (4). Despite major advances in structural biology, obtaining atomic-level insight into protein-membrane interactions under native-like conditions remains challenging (5–7).

At a fundamental level, membrane association is governed by the thermodynamics of partitioning between water and the bilayer. Within this framework, amino acid side-chain insertion free energies are key determinants of membrane protein folding, topology, orientation, and stability (5, 8). Experimental studies of transmembrane helix insertion established hydrophobicity scales that relate side-chain energetics to partitioning within the bilayer (9, 10). Complementary molecular dynamics (MD) simulations have quantified potentials of mean force (PMFs) for side-chain insertion and revealed consistent trends: hydrophobic residues favor the bilayer core, polar and charged residues are excluded from it, and aromatic residues often stabilize interfacial regions (11–13). These thermodynamic preferences are consistent with the compositional biases observed in membrane protein structures (14, 15).

Computationally, insertion PMFs are most often determined by umbrella sampling along the bilayer normal (16, 17). Although rigorous, this approach requires biasing potentials, careful overlap between windows, and substantial equilibration. These requirements increase computational cost and complicate systematic comparisons across multiple residue types, protonation states, and lipid bilayer compositions.

Here, we present an alternative framework for computing side-chain insertion energetics from unbiased MD simulations performed at controlled solute concentrations. Equilibrium depth distributions are sampled directly and converted into PMFs by Boltzmann inversion. This approach avoids biasing restraints while preserving bilayer structure, enabling efficient and internally consistent determination of insertion free energies. We apply this approach to 28 amino acid side-chain analogs, including multiple protonation states, in a 1-palmitoyl-2-oleoyl-*sn*-glycero-3-phosphocholine (POPC) bilayer. The resulting free-energy profiles are validated against experimentally derived hydrophobicity scales. As a consistent reference for systematic comparisons across residue classes and protonation states, we focus here on a well-characterized single-component POPC bilayer. Extension to other lipid bilayer compositions can be achieved within the same unbiased PMF framework.

## METHODS

The partitioning of amino acid side-chain analogs in a POPC bilayer was investigated by MD simulations. Free-energy profiles along the bilayer normal were obtained by Boltzmann inversion of equilibrium positional distributions.

Amino acid side-chain analogs were constructed following established protocols (11, 13, 18). Side chains were truncated at the *β*-carbon, the *α*-carbon was replaced by a proton, and the partial charge of the *β*-carbon was adjusted accordingly. Multiple protonation states were considered where applicable. Glycine was represented by diglycine (GLYD) to capture backbone partitioning, whereas the PRO analog retained the backbone NH group and the *α*-carbon. In total, 28 analogs were simulated, and their chemical structures are shown in Fig. 1. Force field topologies and parameters were taken from the CHARMM36m all-atom additive protein force field (19, 20). Missing parameters for the PRO, GLU^0^, HSE and HSP^+^ analogs were assigned by analogy to existing CHARMM parameters. Topology and parameter files, system input files, PDB files, analysis scripts, and extracted datasets generated in this study are available in a public GitHub repository (21).

**Figure 1:**
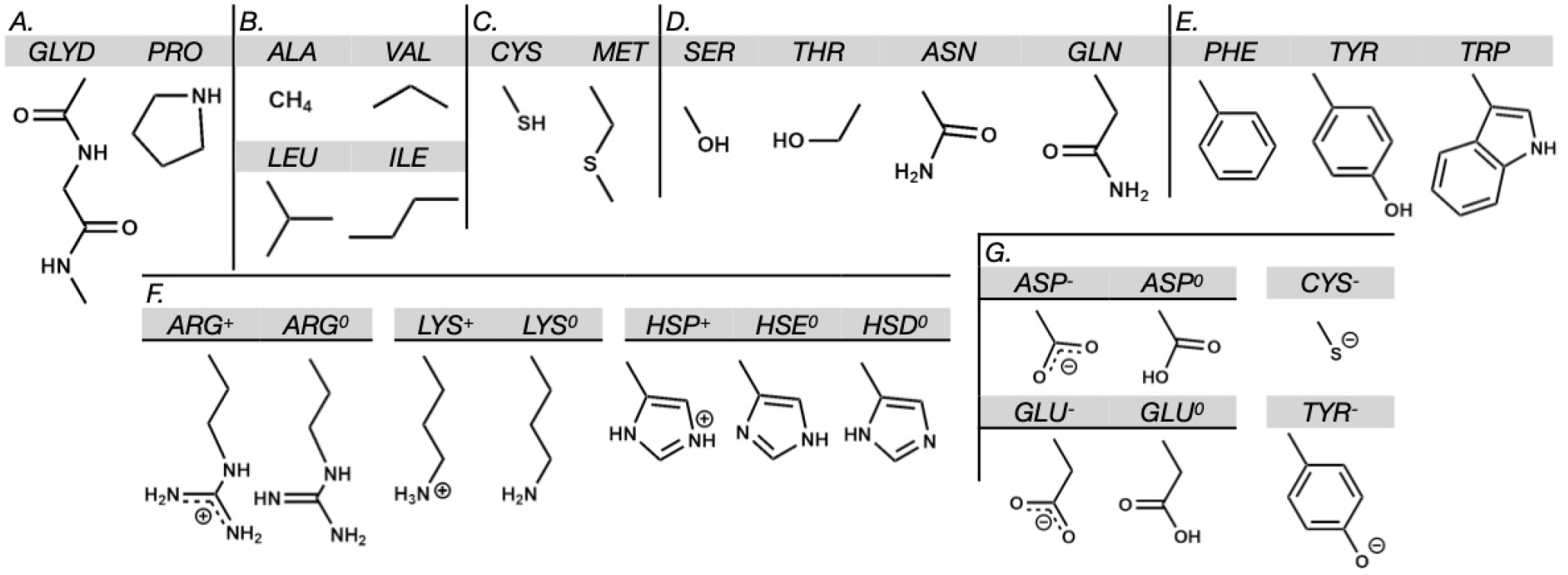
Chemical structures of the 28 analogs, labeled with the three-letter amino acid codes (gray background) for ease of reference, except diglycine, designated GLYD. The set includes 19 amino acid analogs, 8 alternative protonation states, and 1 backbone-related model. Analogs are grouped into seven categories: (A) backbone-related, including proline; (B) hydrophobic; (C) sulfur-containing; (D) polar uncharged; (E) aromatic; (F) positively charged and their neutral or deprotonated forms (ARG^+^, LYS^+^, HSE^0^, HSD^0^ as standard states); and (G) negatively charged and their neutral or protonated forms (ASP^−^, GLU^−^, CYS^0^, TYR^0^ as standard states).

A POPC bilayer was constructed using the Membrane Builder module of CHARMM-GUI (version 3.7) (22–28). The system contained 32 POPC lipids per leaflet and was solvated with water layers 50 Å thick on each side. Sodium and chloride ions were added to neutralize the system and reach a final salt concentration of 150 mM. The final box dimensions were 46.75 × 46.75 × 145 Å^3^, for a total of approximately 28,000 atoms.

For each of the 28 analogs (Fig. 1), an independent system was generated. In each system, 26 analog molecules were initially placed at random in the aqueous phase with Packmol (29) and VMD-based scripts (30), corresponding to a nominal concentration of 200 mM. This concentration was chosen to balance sampling efficiency with limited perturbation of bilayer structure, as assessed from area per lipid, bilayer thickness, and acyl-chain order parameters relative to a bilayer-only control (Supporting Material, Section S1). Overlapping water molecules were removed, and ion counts were adjusted to maintain charge neutrality. In total, 29 systems were generated: 28 containing one analog species and one bilayer-only control.

### Simulation details

MD simulations were performed with NAMD 2.14 (31). Amino acid analogs were modeled with the CHARMM36m force field (19, 20), lipids with CHARMM36 (32, 33), and water with the TIP3P model (34). Simulations were carried out in the isothermal–isobaric (NPT) ensemble with periodic boundary conditions and a 2 fs integration time step. Temperature was maintained at 303.15 K using Langevin dynamics with a damping coefficient of 1 ps^−1^, and pressure was maintained at 1 atm using a Nosé–Hoover Langevin piston.

Bond lengths involving hydrogen atoms were constrained using SETTLE for water molecules (35) and SHAKE for all other bonds involving hydrogens (36). Short-range electrostatic and Lennard-Jones interactions were truncated at 12 Å, with a switching function applied to the Lennard-Jones potential between 10 and 12 Å. Long-range electrostatics were treated using the particle mesh Ewald (PME) method (37), using sixth-order interpolation and a maximum grid spacing of 1.0 Å. PME calculations were performed at every integration step, and the nonbonded pair list was updated every 10 steps. Trajectory frames were saved every 10 ps. For each system, three independent 1,000 ns simulations were performed using different initial velocity seeds.

### Trajectory analysis

Analyses were performed on the final 600 ns of each trajectory, yielding 1800 ns of analyzed data per system. Unless otherwise noted, PMFs were computed from the full analog population. Some analogs occasionally formed intermolecular contacts, defined by at least one inter-analog heavy-atom distance shorter than 4.5 Å. For analogs showing substantial associated populations, PMFs were also computed separately for monomeric and full populations. The fraction of each analog observed in the monomeric state is reported in Table S2 (Section S2).

Prior to analysis, trajectories were recentered along the bilayer normal (*z* axis) by placing the center of mass of the bilayer at the origin. Owing to bilayer symmetry, properties were averaged over both leaflets. Mean distributions and associated standard errors were estimated from nine 200-ns block averages derived from the final 600 ns of the three independent simulations. Bilayer density profiles were calculated with the VMD Density Profile tool (38), whereas analog center-of-mass distributions along *z* were computed with MDAnalysis (39, 40).

The orientation of aromatic analogs (PHE, TRP, and TYR) within the bilayer was quantified using two angular descriptors: *θ*_1_, defined as the angle between the normal vector to the aromatic ring plane and the bilayer normal, and *θ*_2_, an in-plane angle defined between two reference atoms on the aromatic ring. For PHE and TYR, *θ*_2_ was defined using atoms *CG* and *CZ*; for TRP, using *CZ*3 and *CE*2. A schematic illustration of these geometric definitions is provided in Fig. S8.

### PMF calculations

For each analog, a potential of mean force (PMF) along the bilayer normal was computed from the equilibrium positional probability distribution *g*(*z*) using Boltzmann inversion:

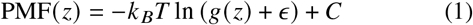

where *k* _*B*_ is the Boltzmann constant, *T* is the absolute temperature, *ϵ* is a small constant (10^−5^) introduced to prevent numerical instabilities when *g* (*z*) approaches zero, and *C* defines the zero of free energy. The probability istribution *g* (*z*) was computed as a normalized histogram of the analog center-of-mass *z* coordinate using 1.0 Å bins:

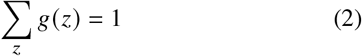

The constant *C* was chosen so that the PMF averaged over the bulk-water region (40 ≤ *z* ≤ 50 Å) was zero.

### pK_a_ calculations

Depth-dependent pK_a_ profiles were derived from the free-energy difference between the charged and neutral forms of each titratable analog. The free-energy difference at depth *z* was defined as:

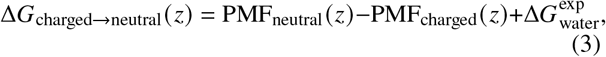

where both PMFs were referenced to zero in bulk water and 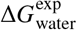 is the experimental free-energy cost of protonation or neutralization in water, derived from experimental aqueous pK_a_ values (41).

The pK_a_ at depth *z* was calculated as:

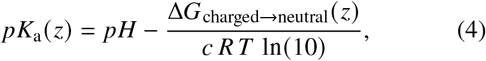

where *c* = +1 for the deprotonation of positively charged forms and *c* = −1 for the protonation of negatively charged forms, *R* is the gas constant, and *T* is the temperature. The reference pH was set to 7.0. Depth ranges in which the charged analog showed insufficient sampling were excluded. This formulation recovers the experimental bulk pK_a_ at large *z* values (Fig. 5, *z* > 40 Å).

## RESULTS

PMF profiles were interpreted using the four-region bilayer model introduced by Marrink et al. (42) and widely applied to membrane insertion studies (11, 13). As illustrated in Fig. 2, Region I corresponds to the hydrophobic core (*z* < 9.5 Å), Regions II and III define the inner and outer interfacial layers (9.5–30 Å), and Region IV represents bulk water (*z* > 30 Å). This regional partitioning is used throughout the Results to describe insertion barriers, preferred depths, and free-energy minima.

**Figure 2:**
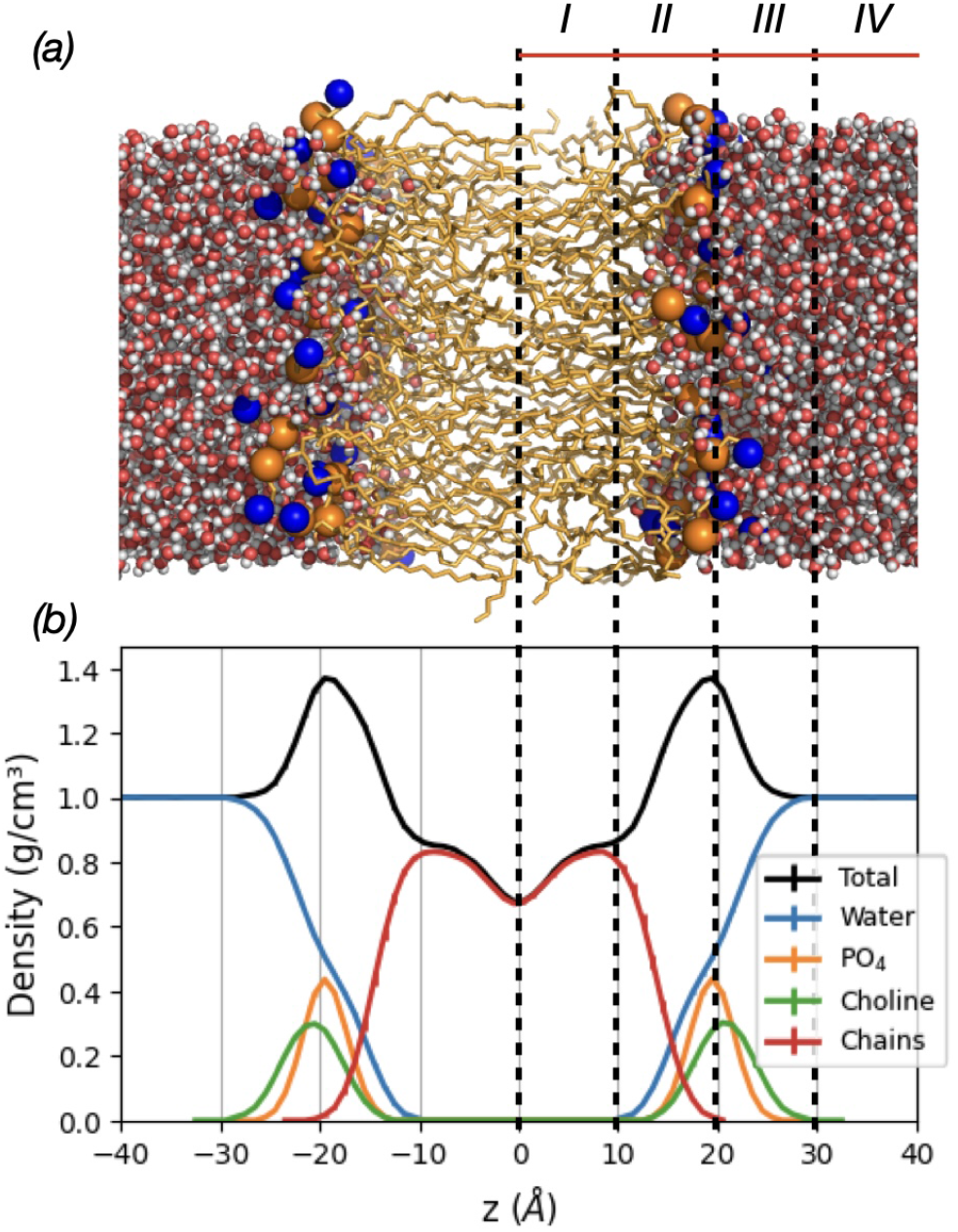
(a) Snapshot of a POPC bilayer. Water molecules are shown as red (oxygen) and white (hydrogen) spheres; phosphorus and nitrogen atoms are shown in orange and blue, respectively; hydrocarbon tails are displayed as orange sticks. (b) Density profiles of selected bilayer components along the bilayer normal (*z*-axis). The bilayer is partitioned into four regions (I to IV).

### Aliphatic and sulfur-containing analogs

The PMF profiles for aliphatic side-chain analogs (ALA, VAL, LEU, and ILE) exhibited shallow interfacial barriers in Region III and progressively favorable insertion toward the bilayer center (Fig. 3). VAL, LEU, and ILE displayed nearly identical profiles characterized by deep minima within Region I. In contrast, ALA exhibited a markedly shallower minimum, consistent with its lower hydrophobicity.

**Figure 3:**
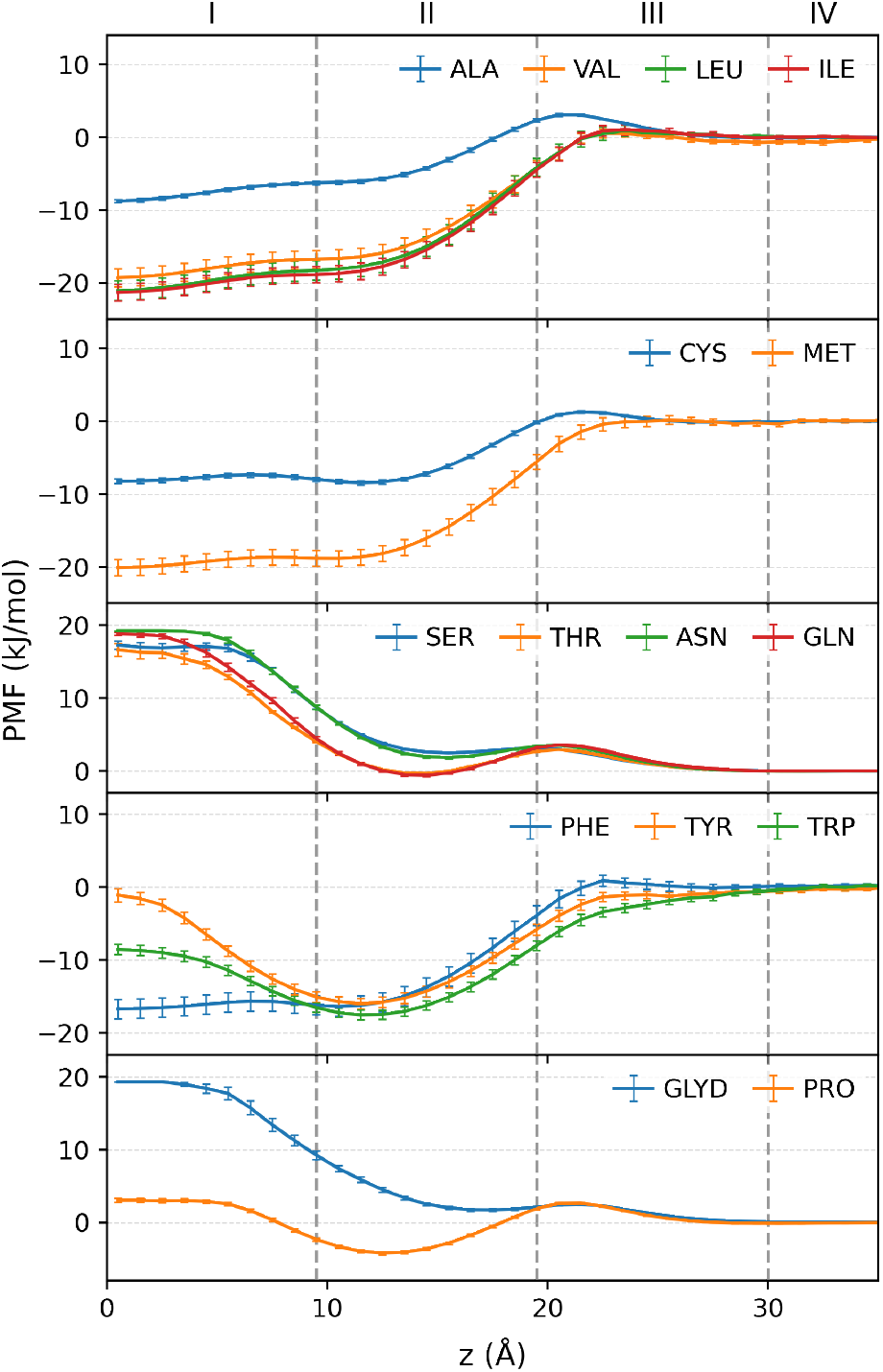
PMF profiles of amino acid analogs. Rows correspond to aliphatic, sulfur-containing, polar, aromatic, and backbone-related analogs. Free energies are referenced to zero in bulk water. Error bars show standard errors estimated from nine 200-ns block averages across three independent simulations.

CYS and MET showed similar overall trends (Fig. 3). Both favored insertion toward the bilayer center, with MET displaying a deep minimum comparable to the larger aliphatic residues, whereas neutral CYS showed higher free energies and a shallower minimum. In contrast, the deprotonated form of CYS exhibited a steep free-energy increase from the solvent-facing interface toward the bilayer center (Fig. 4), reflecting the exclusion of charged species from the hydrophobic interior.

**Figure 4:**
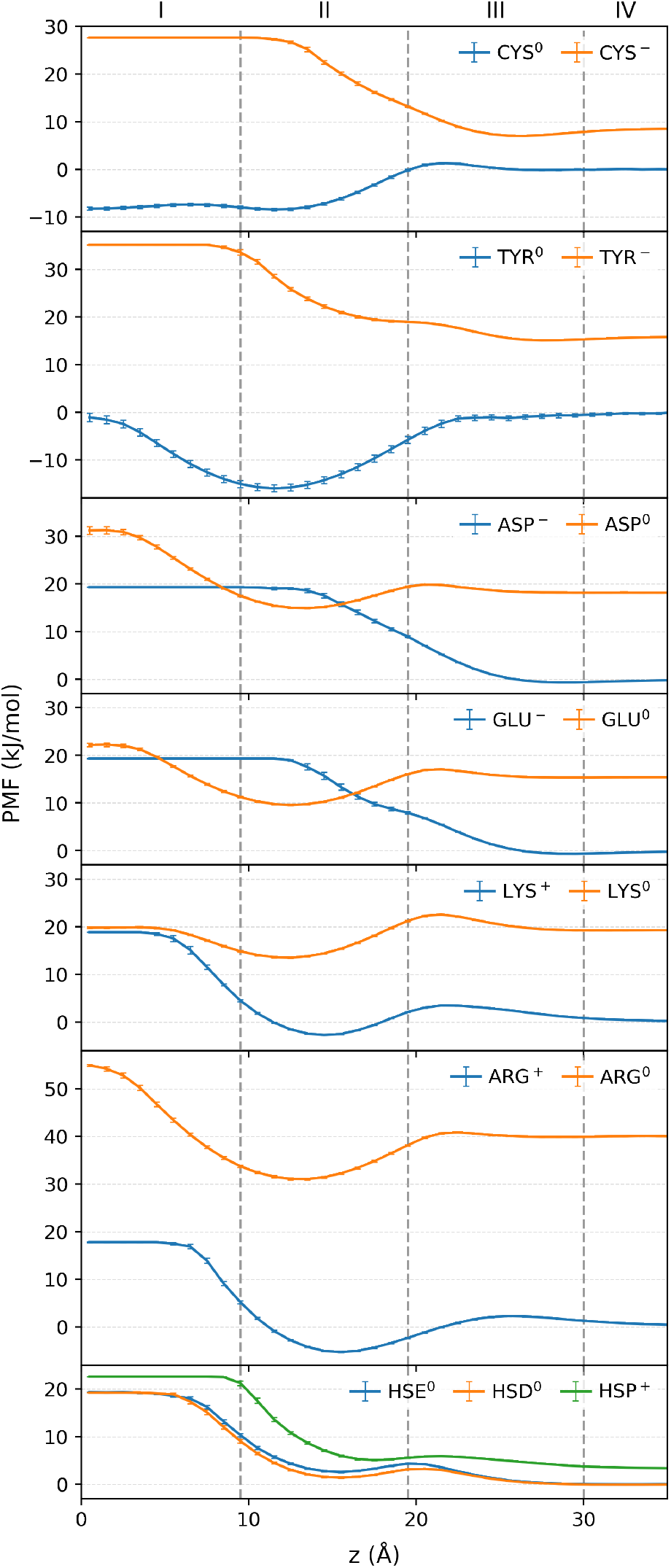
PMF profiles of analogs with different protonation states. For protonation states that are non-standard in bulk water at pH 7, PMFs were shifted by the free-energy cost of protonation or neutralization derived from experimental pK_a_ values (41). Error bars show standard errors estimated from nine 200-ns block averages across three independent simulations.

### Polar and backbone-related analogs

Polar analogs (ASN, GLN, SER, and THR) displayed unfavorable insertion across the bilayer (Fig. 3). All four showed modest interfacial barriers near the boundary between Regions II and III. ASN and SER remained unfavorable at all depths, whereas GLN and THR displayed near-neutral free energies in Region II before becoming strongly unfavorable toward the bilayer center. The backbone-related analogs followed the same overall trend. GLYD exhibited a polar-like profile with unfavorable insertion across the bilayer, whereas the PRO analog displayed a shallow interfacial minimum followed by a progressive increase in free energy toward the bilayer center, where insertion became unfavorable.

### Aromatic analogs

Aromatic analogs (PHE, TYR, and TRP) exhibited favorable insertion profiles (Fig. 3). All three displayed free-energy minima within Region II, with TRP exhibiting a slightly deeper minimum than PHE and TYR. Beyond the interfacial minimum, the free energies of TYR and TRP increased toward the bilayer center, indicating reduced stability in the hydrophobic core. In contrast, the PMF of PHE remained relatively flat beyond its minimum, maintaining comparable free energies from Region II to the bilayer center. TYR and TRP reached their minima without a detectable interfacial barrier, whereas PHE exhibited a small interfacial barrier prior to insertion. The deprotonated form of TYR (Fig. 4) showed a steep increase in free energy upon insertion toward the bilayer core, becoming strongly unfavorable and reaching the upper limit of the PMF range.

### Charged and ionizable analogs

PMF profiles for acidic (ASP, GLU), basic (ARG, LYS), and ionizable (HIS) analogs are shown in Fig. 4.

#### Acidic residues

Anionic ASP and GLU exhibited rapidly increasing free energies upon entering Region III, with no favorable insertion at any bilayer depth. Their neutral forms displayed small interfacial barriers and shallow minima within Region II, followed by progressively unfavorable insertion toward the bilayer center. However, when accounting for the free-energy cost of protonation, these minima do not correspond to thermodynamically favorable insertion relative to bulk water.

#### Basic residues

ARG and LYS, in both charged and neutral forms, exhibited modest interfacial barriers followed by shallow minima within Region II. Deeper insertion resulted in a steep rise in free energy, reaching the upper limit of the PMF scale. Only the cationic forms exhibited weak interfacial stabilization.

#### Ionizable aromatics

All histidine analogs (HSE, HSD, and HSP) exhibited unfavorable insertion, characterized by a small barrier near the interface and a steep rise toward the hydrophobic core.

### Protonation effects

Depth-dependent pKa profiles were derived from free-energy differences between charged and neutral states according to Eq. 4 and are shown in Fig. 5. Because charged analogs only rarely sampled the bilayer core, Region I was excluded from the pKa analysis.

**Figure 5:**
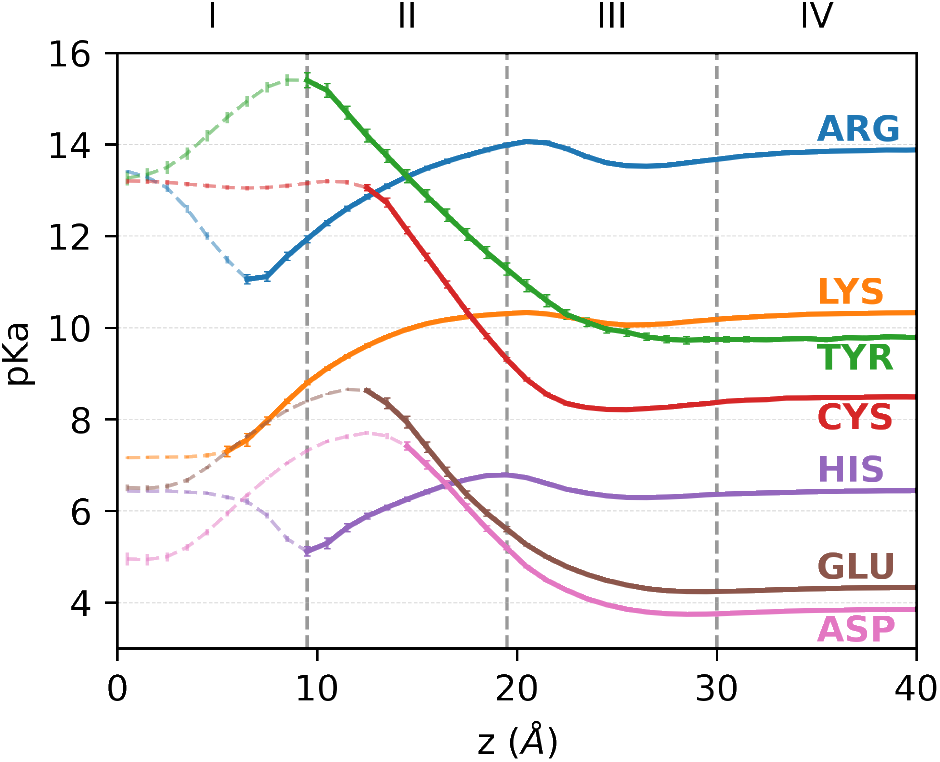
Depth-dependent side-chain pK_a_ profiles across the POPC bilayer. pK_a_ values were calculated from free-energy differences between charged and neutral forms using Eq. 4. Experimental aqueous pK_a_ values were taken from Platzer *et al*. (41). Error bars represent propagated standard errors derived from the PMFs of the charged and neutral analogs.

Bilayer-induced desolvation strongly favored neutral forms, with pronounced pKa shifts emerging at the interface between Regions II and III. Upon insertion into Region II, LYS and ARG exhibited substantial downward pKa shifts, reflecting destabilization of their protonated states, whereas ASP, GLU, CYS, and TYR showed upward pKa shifts indicative of stabilization of the protonated forms. HIS displayed comparatively modest variations and remained close to its aqueous pKa across the bilayer.

### Aromatic-ring orientations

Aromatic-ring orientations were quantified using angular descriptors defined in Materials and Methods and analyzed as a function of bilayer depth. Near the boundary between Regions I and II, TYR and TRP exhibited well-defined preferred orientations, whereas PHE displayed broader angular distributions, indicating weaker orientational preferences (Fig. 6). Across all three aromatic analogs, orientations with the ring plane aligned along the bilayer normal were favored. Representative orientations are shown in Fig. S9.

**Figure 6:**
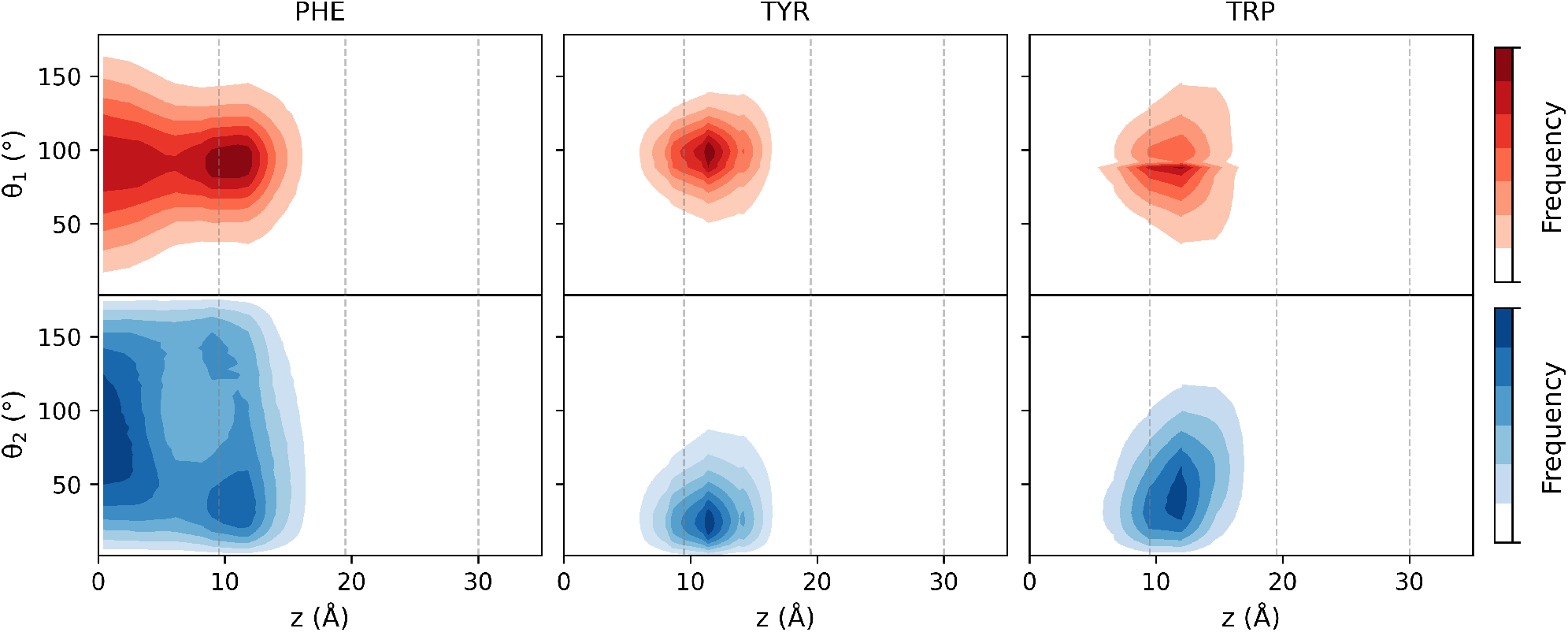
Depth-resolved angular distributions for aromatic analogs. Red probability maps correspond to *θ*_1_, the angle between ring normal and bilayer normal. Blue probability maps correspond to *θ*_2_, the angle between the long axis of the ring and the bilayer normal. Angle definitions are illustrated in Fig. S8. The frequencies in these probability maps are normalized. Results for TYR^−^ are not shown due to insufficient sampling at bilayer depths relevant for orientational analysis.

### Association of analogs

A limited subset of analogs sampled substantial associated populations during the simulations. The fraction of each analog observed in the monomeric state is reported in Table S2, and PMF profiles for monomeric and full populations are shown in Fig. S7 for analogs with associated fractions exceeding 40%. These analogs include VAL, LEU, ILE, MET, PHE, TYR^0^, TRP, and ARG^0^. With the exception of neutral ARG, all correspond to hydrophobic or aromatic residues. Comparison of monomeric and full-population PMFs indicates only slight changes in the insertion free-energy profiles. Additional simulations of selected residues performed at half the concentration (100 mM) using the same protocol produced no substantial changes in the PMF profiles, supporting the absence of strong cooperative effects.

### Hydrophobicity-scale comparison

Comparison of the calculated insertion free energies with experimental hydrophobicity scales for residue partitioning into POPC and DLPC bilayers (9, 10) showed strong agreement across residue classes, with correlation coefficients of 0.85–0.87 (Fig. S10). Hydrophobic analogs clustered in the low–free-energy regime, whereas polar residues populated the high–free-energy region, consistent with experimental trends. The full comparison, including individual residue contributions and both experimental scales, is provided in the Supporting Material (Section S4).

## DISCUSSION

In this study, we systematically quantified bilayer insertion free energies of 28 amino acid side-chain analogs in a POPC bilayer by deriving PMFs from depth distributions sampled in unbiased MD simulations. This strategy provides an efficient alternative to umbrella sampling (11, 13) and free-energy perturbation approaches (12, 13), enabling broad and internally consistent comparisons using a single simulation protocol and reference state across all residues and protonation states.

### Methodological considerations

Boltzmann inversion of depth distributions provides a simple and computationally efficient route for estimating membrane insertion PMFs. By eliminating the need for biasing potentials, this approach avoids challenges associated with window overlap, force-constant tuning, and equilibration that accompany umbrella sampling. Moreover, when solute concentrations are chosen appropriately, the method preserves intrinsic bilayer properties (Section S1, Figs. S1-S3 and S5-S6) while achieving extensive configurational sampling at modest computational cost.

Several limitations should nonetheless be acknowledged. First, side-chain analogs were truncated at C_*β*_ and therefore do not explicitly capture backbone-mediated desolvation. This contribution was quantified separately using the diglycine analog, allowing us to isolate and assess the backbone desolvation penalty.

Second, although intermolecular association was observed for a limited number of primarily hydrophobic and aromatic analogs, its effect on the PMFs was small. For residues with substantial associated populations, PMFs computed from monomeric and full populations differed only slightly. In addition, simulations repeated at half the solute concentration produced no significant changes in the PMF profiles. Together, these observations argue against strong cooperative effects and indicate that association contributes only marginally to the insertion energetics under the conditions studied. This interpretation is consistent with previous computational studies showing that side-chain association in membrane environments makes limited, context-dependent energetic contributions (14, 18).

Third, sampling efficiency was intrinsically reduced in high-free-energy regions, such as the bilayer center for polar or charged analogs and bulk water for hydrophobic analogs. This limitation resulted in increased statistical uncertainty or PMF plateaus and reflects a well-recognized constraint of unbiased MD approaches to PMF estimation (16, 17).

Finally, the simulations employed additive CHARMM36 and CHARMM36m force fields. While polarizable lipid models can more accurately capture bilayer dipole potentials and permeation energetics (43, 44), they remain computationally demanding and do not uniformly yield improved bilayer properties (45). In addition, polarizable parameters are not yet available for the full diversity of lipid species found in biological membranes, limiting their applicability for systematic comparisons across multiple membrane compositions, an important direction for future extensions of this work.

### General features of insertion PMFs across bilayer regions

The PMFs reflect the heterogeneous physicochemical environment of the bilayer, spanning the bulk aqueous phase to the hydrocarbon core (Figs. 3 and 4). As expected, hydrophilic residues remain energetically favored in the aqueous phase, whereas hydrophobic aliphatic residues preferentially partition into the bilayer interior. Most analogs exhibited a characteristic interfacial signature: a shallow barrier at the aqueous interface (Region III), followed by modest stabilization within the inner headgroup region (Region II). This behavior reflects the transition from the hydrated, densely packed phosphate/choline environment to the increasingly nonpolar, yet still structured, upper acyl-chain region. Similar interfacial stabilization has been reported in previous amino acid insertion studies (11, 13) and is a well-established feature of amphiphilic drugs and small molecules (46).

#### Hydrophobic and sulfur-containing residues

Hydrophobic residues exhibited increasingly favorable insertion with side-chain size (Fig. 3), albeit with non-monotonic trends among branched aliphatic residues. Insertion free energies ranged from approximately −9 kJ/mol for ALA to −21 kJ/mol for ILE, with LEU exhibiting insertion energetics closer to VAL than to ILE. These trends are consistent with previous computational studies (11, 13) and slightly less favorable than hexane partitioning (*∼* −25 kJ/mol) (47).

Sulfur-containing analogs exhibited intermediate behavior. MET displayed an insertion profile similar to those of the larger aliphatic residues, whereas CYS showed weaker stabilization, consistent with its modest polarity. These results differ from those of MacCallum *et al*. (11), who reported nearly identical profiles for MET and CYS. The discrepancy likely reflects differences in sulfur partial charge assignments: sulfur carries substantially more negative charge in OPLS-AA (48, 49) than in CHARMM36, increasing desolvation penalties. This interpretation is supported by the close agreement between our CYS PMF and that reported by Pogorelov *et al*. (13), who also employed a CHARMM-based force field, albeit using an HMMM membrane model and umbrella sampling. Consistent with this view, our depth-dependent behavior more closely resemble those observed in MemProtMD simulations (50), although it should be noted that CYS and MET are rarely solvent-exposed in membrane proteins (51).

#### Aromatic residues

Aromatic residues exemplify the interplay between hydrophobicity, polarity, and orientational preferences at bilayer interfaces. PHE exhibited a hydrophobic insertion profile with a nearly flat free-energy landscape across the bilayer core and a midplane minimum of approximately −16 kJ/mol, consistent with previous work (11, 13). In contrast, TYR and TRP displayed stabilization within the inner headgroup region (Region II), with minima of approximately −15 kJ/mol for TYR and −17 kJ/mol for TRP. Beyond the interfacial region, insertion free energies increased toward the bilayer center, reaching near-neutral values for TYR, whereas TRP remained modestly stabilized, with a midplane free energy of approximately −8 kJ/mol.

Comparisons with previous studies highlight the sensitivity of aromatic insertion energetics to force-field choice and bilayer composition. Overall trends are similar to those reported by MacCallum *et al*. and Pogorelov *et al*., although quantitative differences are observed. In particular, both studies reported unfavorable insertion of tyrosine at the bilayer center, whereas our results indicate near-neutral energetics at this depth. These differences likely reflect variations in midplane polarity and in partial charge assignments on aromatic heteroatoms.

Despite quantitative differences, our results are consistent with experimental and computational evidence showing that simple aromatic compounds such as benzene partitions uniformly across POPC bilayers (52), whereas indole, TRP, and TYR preferentially localize at the hydrocarbon-polar interface (53–56). Orientation analyses support this interpretation: TYR and TRP preferentially adopted orientations in which the ring plane was aligned along the bilayer normal at interfacial depths (Region II), facilitating stabilization through interactions between polar substituents and lipid headgroups (Fig. 6). PHE sampled broader orientations while retaining a weak preference for similar alignment.

### Polar and charged residues

Polar and charged residues exhibited insertion behaviors distinct from those of aromatic analogs. SER, ASN, ASP, GLU, and HIS were unfavorable at all bilayer depths, whereas THR and GLN approached near-neutral free energies within Region II. LYS and ARG showed modest interfacial stabilization, reflecting favorable hydration and transient headgroup interactions.

Overall insertion trends for SER, ASN, ASP, and ARG are consistent with those reported by Pogorelov *et al*. (13), although higher free-energy plateaus were observed near the bilayer center in our simulations. In contrast, MacCallum *et al*. (11) reported systematically deeper interfacial minima for polar and charged residues, resulting in more favorable insertion in Region II. These quantitative differences are most plausibly attributed to differences in force-field parametrization rather than methodological approach.

Unlike several previous umbrella-sampling studies (11, 13, 57), we did not observe persistent water defects accompanying the insertion of polar or charged analogs. Because our simulations were unbiased, solutes were free to diffuse naturally rather than being restrained at unfavorable depths. This suggests that water defects reported in biased simulations may, in part, be favored by restraint at unfavorable depths or by incomplete equilibration, although such defects can arise and be stabilized in peptide or protein contexts (58).

Depth-dependent pK_a_ profiles (Fig. 5) derived from the PMFs revealed substantial shifts for all titratable residues. Basic residues exhibited pronounced downward pK_a_ shifts upon insertion into Region II, whereas ASP, GLU, CYS, and TYR showed marked upward shifts stabilizing their protonated forms. These trends are consistent with previous computational and theoretical studies (11, 59, 60) and underscore the strong coupling between protonation state, insertion depth, and bilayer polarity, key determinants of peptide stability and function in membrane environments.

### Backbone-related analogs: GLYD and PRO

The backbone-containing analog GLYD and the cyclic PRO analog illustrate how backbone polarity and conformational constraints modulate bilayer insertion energetics. GLYD exhibited a uniformly unfavorable insertion profile, with minimal interfacial stabilization and a steep free-energy increase toward the bilayer center. This behavior is consistent with experimental and computational studies showing that unshielded peptide backbones incur large desolvation penalties unless compensated by hydrogen bonding or secondary structure formation (5, 8, 9, 58). In contrast, the PRO side-chain analog displayed a shallow interfacial minimum and near-neutral stabilization across the upper acyl-chain region. This behavior reflects the reduced backbone polarity and conformational constraint associated with the cyclic structure of proline. These features facilitate partial penetration into Region II without imposing large energetic penalties. The behavior of PRO has direct implications for membrane protein structure: proline frequently introduces kinks or hinges in transmembrane helices by disrupting backbone hydrogen bonding (61), modulating helix packing, orientation, and dynamics. The near-neutral insertion energetics observed here are consistent with this functional role.

### Interpretation of hydrophobicity comparison

The strong correspondence between our PMFs and experimental hydrophobicity scales, presented in the Supporting Material (Section S4), is notable given differences in effective protonation conditions (e.g., standard protonation states in our simulations versus pH 8.0 for Wimley-White and pH 3.8 for Moon-Fleming), lipid composition (POPC vs. DLPC bilayer), and molecular context (isolated side-chain analogs vs. residues embedded in host-guest peptide or transmembrane proteins) across datasets. This agreement indicates that the dominant physicochemical determinants of membrane partitioning are consistently captured across approaches.

## CONCLUSION

In this study, unbiased molecular dynamics simulations were used to quantify amino acid side-chain insertion free energies in a POPC bilayer. Equilibrium depth distributions of 28 analogs, including multiple protonation states, were converted into potentials of mean force, enabling systematic and internally consistent comparison of residue-specific insertion energetics within a single simulation framework. The resulting PMFs recapitulate the main thermodynamic features of bilayer partitioning, including preferential localization of hydrophobic residues to the bilayer core, interfacial stabilization of aromatic residues, and exclusion of polar and charged species from the hydrophobic interior.

Depth-dependent pK_a_ profiles and orientational analyses further show how desolvation, protonation equilibria, and aromatic-ring alignment modulate insertion energetics across bilayer regions. The strong agreement with experimental hydrophobicity scales and with structural trends observed in membrane proteins supports the robustness of the approach and its ability to capture the dominant thermodynamic determinants of protein-membrane interactions.

Although the present framework has limitations, including truncated analog representations, intermolecular association for a limited subset of analogs, reduced sampling in highfree-energy regions, and the use of additive force fields, it provides an efficient and extensible strategy for characterizing bilayer insertion energetics. Extension to more diverse lipid compositions and more complex peptide or protein contexts should broaden the applicability of this framework and further refine our understanding of membrane-associated processes.

## AUTHOR CONTRIBUTIONS

S. Bories designed and performed the research, carried out all simulations, analyzed the data, and drafted the manuscript. P. Lague supervised the research and contributed to writing and revising the manuscript.

## DECLARATION OF GENERATIVE AI AND AI-ASSISTED TECHNOLOGIES IN THE MANUSCRIPT PREPARATION PROCESS

During the preparation of this work, the authors used Chat-GPT (OpenAI) to assist with language editing, wording refinement, and improvement of the clarity and readability of the manuscript. After using this tool, the authors reviewed and edited the content as needed and take full responsibility for the content of the published article.

## ACKNOWLEDGMENTS

This work was supported by the Natural Sciences and Engineering Research Council of Canada (NSERC) Discovery Grant program (RGPIN-2022-04721) and by the Regroupement Québécois de Recherche sur la Fonction, l’Ingénierie et les Applications des Protéines (PROTEO) – Fonds de recherche du Québec (62). Calculations were performed on supercomputers supported by Calcul Québec and the Digital Research Alliance of Canada. These computing resources are funded by the Canada Foundation for Innovation (CFI), the ministère de l’Économie, de l’Innovation et de l’Énergie du Québec (MEIE), and the Fonds de recherche du Québec (FRQ).

## SUPPORTING MATERIAL

The Supporting Material available through BJ Online includes additional analyses supporting the results discussed in this article, including bilayer structural properties, monomeric fractions and monomer-specific PMFs, aromatic ring orientation analyses, and comparisons with experimental hydrophobicity scales. Data files for the depth distributions and associated PMFs, the data used to generate all figures, and additional topology and parameter files used for simulations of PRO, HSE^0^, HSP^+^, and GLU^0^ are available in a public GitHub repository (21). An online supplement to this article can be found by visiting BJ Online at http://www.biophysj.org.

