## Supplemental Material for "Amino Acid Insertion Energetics in a POPC Bilayer from Unbiased Molecular Dynamics"

### Contents

**Section S1 Bilayer structural properties** 3

**Section S2 Monomeric Fractions and Insertion PMFs** 9

**Section S3 Ring orientations** 11

**Section S4 Hydrophobicity scales** 13

### Section S1 Bilayer structural properties

Trajectories were analyzed to evaluate key bilayer properties, including bilayer thickness ($D_{P-P}$), area per lipid ($A_{PL}$), area compressibility modulus ($K_A$), deuterium order parameters ($S_{CD}$), and mass density profiles (MDPs). These properties were averaged over the final 600 ns of each trajectory across three independent simulations and both leaflets. Unless otherwise stated, standard errors (SE) were estimated from nine 200-ns block averages extracted from those 600 ns segments.

**Table S1**: Calculated POPC bilayer properties with associated SEs and comparison with experimental values.

| Parameter | Unit | Simulation | SE | Exp |
|---|---|---|---|---|
| $D_{P-P}$ | Å | 39.12 | 0.03 | 37.6,[a] 39.1[b] |
| $A_{PL}$ | Å$^2$ | 64.61 | 0.05 | 68.3,[a] 64.3[b] |
| $K_A$ | mN/m | 226 | 6 | 180-330[c] |

[a] Kučerka et al. 2006
[b] Kučerka et al. 2011
[c] Binder and Gawrisch 2001 (298 K)

Each property was compared between the POPC-only bilayer and analog-containing systems to evaluate the impact on bilayer structural properties. For profile-based properties such as $S_{CD}$ and MDPs, deviations were quantified using the normalized absolute difference between the discrete profiles:

$$\%\,\text{Difference} = \frac{\sum_i |P_i - A_i|}{\sum_i P_i} \times 100\%,$$

where $P_i$ denotes the POPC-only reference profile evaluated at bin $i$, and $A_i$ the corresponding profile for the analog-containing system. Lower values indicate higher similarity.

**Bilayer thickness ($D_{P-P}$):** $D_{P-P}$ was computed as the distance between the average $z$-positions of lipid phosphate atoms in the two leaflets.

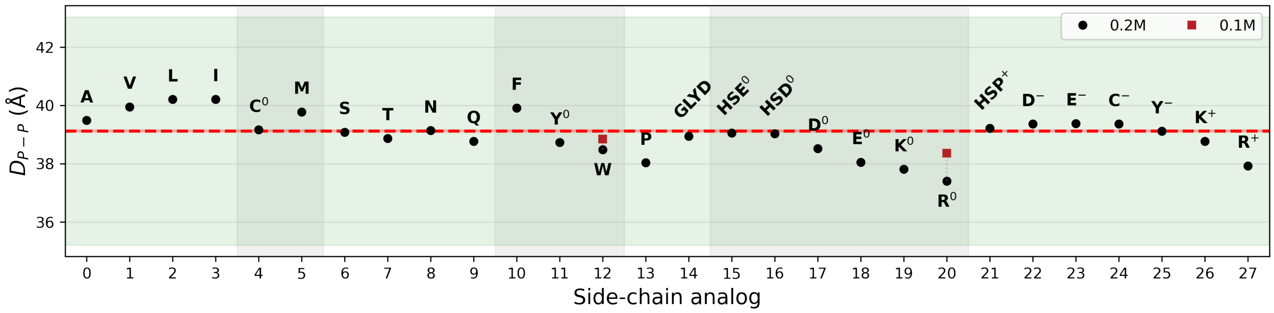

**Figure S1**: Phosphate-to-phosphate bilayer thickness ($D_{P-P}$) for POPC bilayers containing amino acid side-chain analogs, relative to the POPC-only reference. Black symbols show mean values for each system at 0.2 M. Red squares show values from same-length simulations at half the analog concentration (TRP and ARG$^0$; 0.1 M). Means and SEs were estimated from nine 200-ns blocks extracted from the final 600 ns of three independent simulations per system. The red dashed line indicates the POPC-only reference, the green shaded region indicates a $\pm10\%$ reference range, and analogs are grouped by chemical properties. Standard errors are smaller than 0.1 Å.

**Area per lipid ($A_{PL}$):** $A_{PL}$ was computed from the bilayer area in the $xy$ plane ($A_{XY}$) and the total number of POPC lipids ($N_L = 64$) according to

$$A_{PL} = \frac{A_{XY}}{N_L/2}.$$

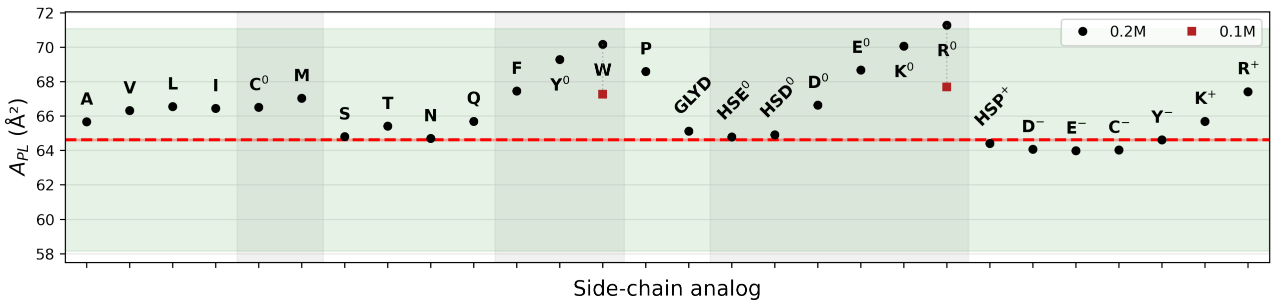

**Figure S2**: Area per lipid ($A_{PL}$) for POPC bilayers containing amino acid side-chain analogs, relative to the POPC-only reference. Plotting conventions are the same as in Fig. S1.

**Area compressibility modulus ($K_A$):** $K_A$, which characterizes the resistance of a bilayer to isotropic area expansion or compression, was computed according to

$$K_A = k_B T \frac{\langle A \rangle}{\langle \delta A^2 \rangle},$$

where $k_B$ is the Boltzmann constant, $T$ the temperature, $\langle A \rangle$ the average lateral area of the simulation box, and $\langle \delta A^2 \rangle$ the mean square fluctuation of $A$.

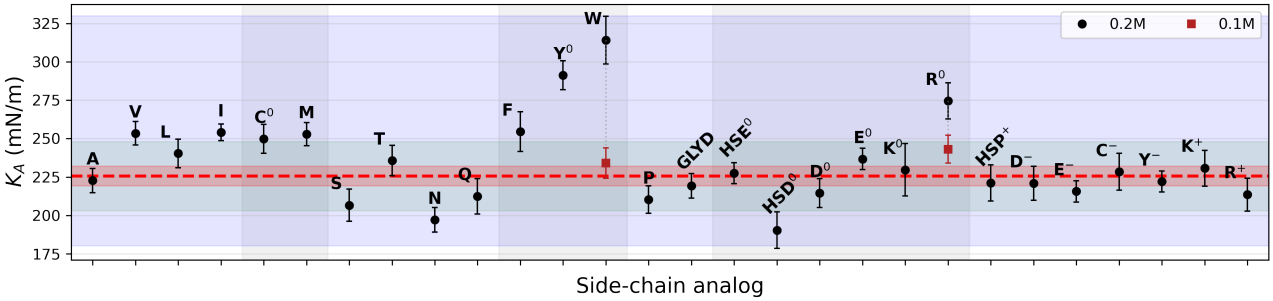

**Figure S3**: Area compressibility modulus ($K_A$) for POPC bilayers containing amino acid side-chain analogs, relative to the POPC-only reference. Plotting conventions are the same as in Fig. S1, except that the blue shaded region indicates the experimental range (180–330 mN/m)[3].

**Deuterium order parameter ($S_{CD}$):** $S_{CD}$, which characterizes acyl chain order, was computed according to

$$S_{CD} = \frac{\langle 3\cos^2\theta - 1\rangle}{2},$$

where $\theta$ is the angle between the C-H bond vector and the bilayer normal ($z$-axis). This parameter was calculated for each carbon atom along both acyl chains of the POPC lipids.

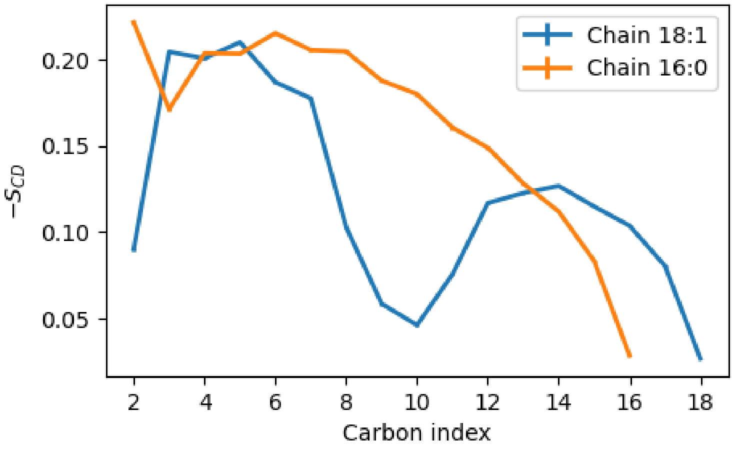

**Figure S4**: Deuterium order parameter ($S_{CD}$) as a function of carbon position along the sn-1 (palmitoyl) and sn-2 (oleoyl) acyl chains in the POPC-only bilayer. The two curves show mean $S_{CD}$ profiles for the sn-1 and sn-2 chains. Error bars show SEs. Means and SEs were obtained from nine 200-ns blocks extracted from the final 600 ns of three independent simulations. Standard errors are smaller than 0.001.

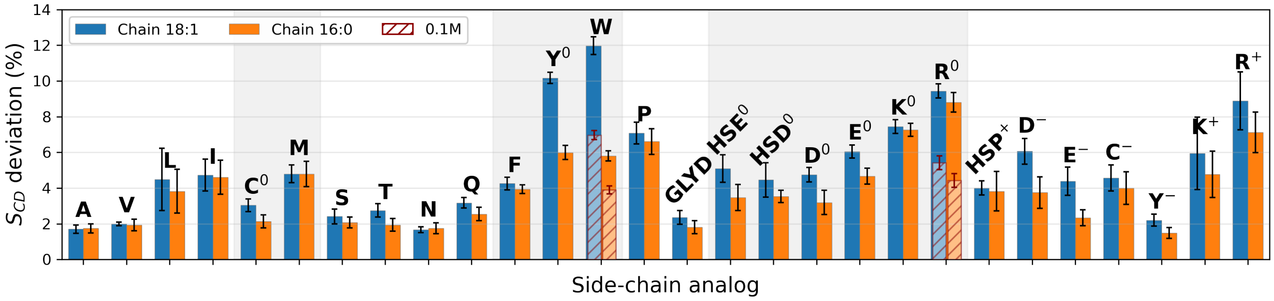

**Figure S5**: Percentage deviation in deuterium order parameter ($S_{CD}$) profiles for POPC acyl chains in bilayers containing amino acid side-chain analogs, relative to the POPC-only reference. Plain bars show mean deviations for each system at 0.2 M, and red hatched bars show values from same-length simulations at half the analog concentration (TRP and $ARG^0$; 0.1 M). Mean values and SE estimates were obtained from nine paired 200-ns block differences between the POPC-only and analog-containing systems, using blocks from the final 600 ns of each of three independent simulations.

**Mass density profiles (MDPs):** Mass density profiles (MDPs) were computed along the $z$-axis for the following components: lipid acyl tails, choline groups, phosphate groups, water, and the total system. Profiles represent averages over the final 600 ns of three independent simulations. The bilayer was centered at $z = 0$. Profiles were averaged over both leaflets.

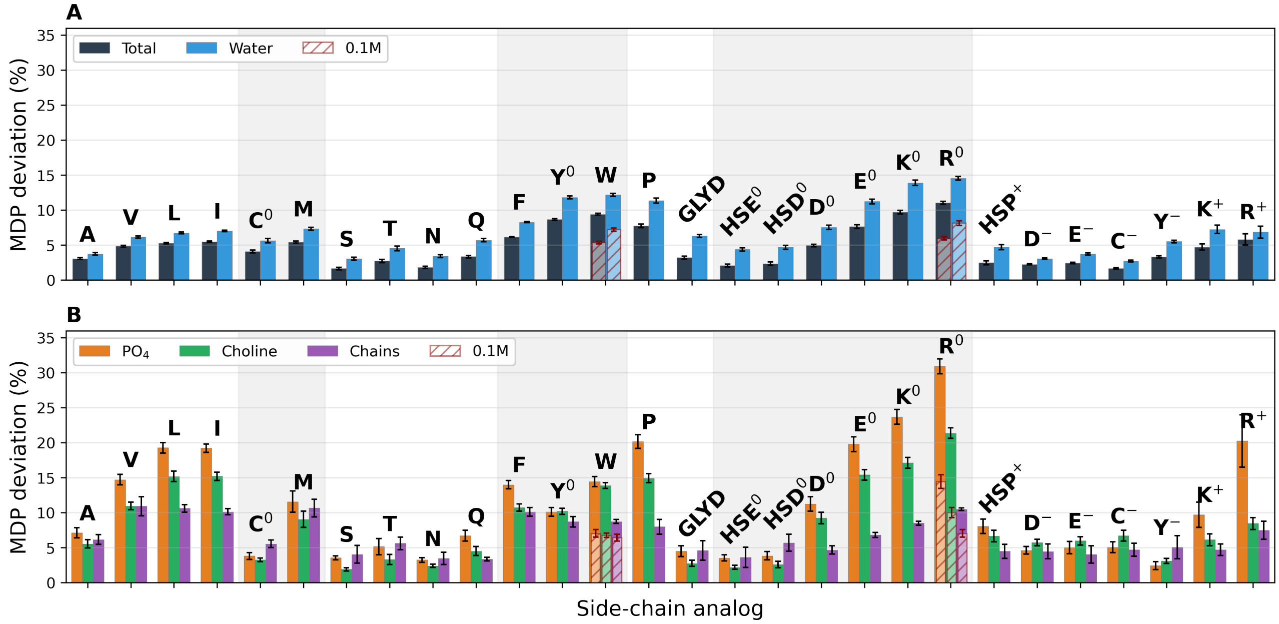

**Figure S6**: Percentage deviation in mass density profiles (MDPs) for POPC bilayers containing amino acid side-chain analogs, relative to the POPC-only reference. Panel A reports total mass density and water, and panel B reports phosphate groups, choline groups, and acyl chains. Plain bars show mean deviations for each system at 0.2 M, and red hatched bars show values from same-length simulations at half the analog concentration (TRP and $ARG^0$; 0.1 M). Mean values and SE estimates were obtained from nine paired 200-ns block differences between the POPC-only and analog-containing systems, using blocks from the final 600 ns of each of three independent simulations.

### Section S2 Monomeric Fractions and Insertion PMFs

**Table S2**: Fraction of each analog observed in the monomeric state during the simulations. Monomers were defined as molecules with no inter-analog heavy-atom contacts shorter than 4.5 Å. Means and standard deviations (SDs) were calculated from nine 200-ns blocks, obtained from the final 600 ns of three independent simulations for each system.

| Analog | Monomer Fraction (%) |
|---|---|
| ALA | $82.6 \pm 1.2$ |
| VAL | $59.8 \pm 0.8$ |
| LEU | $52.1 \pm 0.9$ |
| ILE | $52.7 \pm 0.8$ |
| MET | $54.0 \pm 0.9$ |
| $CYS^0$ | $76.9 \pm 1.4$ |
| SER | $93.3 \pm 0.1$ |
| THR | $90.6 \pm 0.3$ |
| ASN | $87.8 \pm 0.3$ |
| GLN | $84.3 \pm 0.4$ |
| PHE | $52.4 \pm 1.5$ |
| $TYR^0$ | $50.8 \pm 1.9$ |
| TRP | $41.8 \pm 0.8$ |
| PRO | $83.9 \pm 1.5$ |
| GLYD | $68.0 \pm 0.8$ |
| $ASP^0$ | $87.2 \pm 0.5$ |
| $GLU^0$ | $76.2 \pm 2.3$ |
| $HSE^0$ | $85.1 \pm 0.3$ |
| $HSD^0$ | $84.4 \pm 0.3$ |
| $LYS^0$ | $76.8 \pm 0.3$ |
| $ARG^0$ | $56.5 \pm 2.6$ |
| $ASP^-$ | $94.8 \pm 0.1$ |
| $GLU^-$ | $93.2 \pm 0.1$ |
| $CYS^-$ | $97.7 \pm 0.1$ |
| $TYR^-$ | $87.6 \pm 0.3$ |
| $HSP^+$ | $95.1 \pm 0.1$ |
| $LYS^+$ | $93.0 \pm 0.3$ |
| $ARG^+$ | $79.1 \pm 2.3$ |

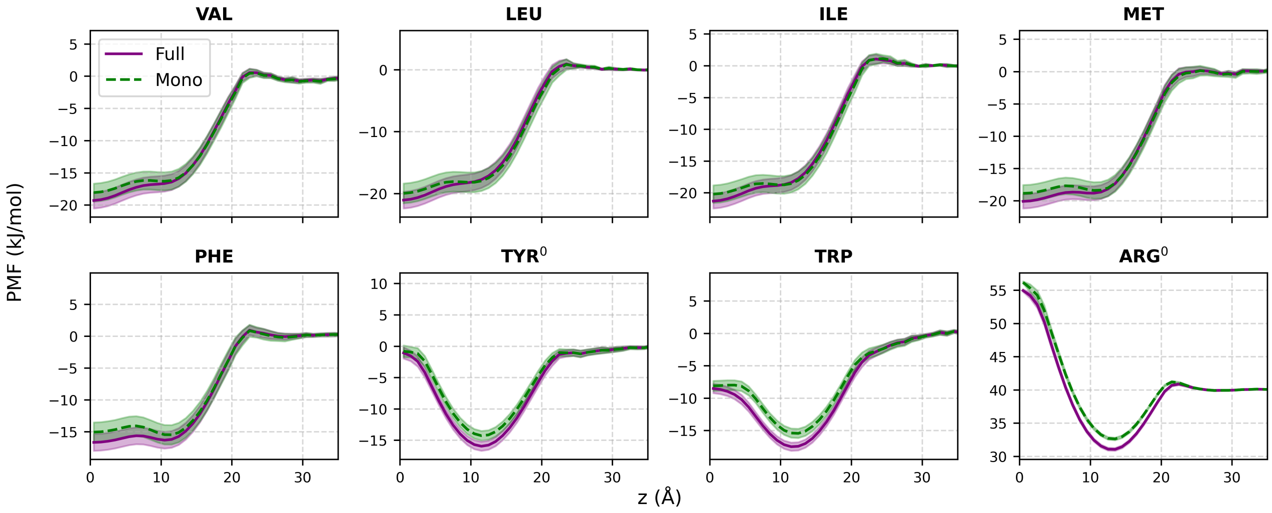

**Figure S7**: Potentials of mean force (PMFs) along the bilayer normal for monomeric and full populations of side-chain analogs showing more than 40% associated species in the POPC bilayer. Green curves show monomeric populations and purple curves show full populations. Monomers were defined as molecules with no inter-analog heavy-atom contacts shorter than 4.5 Å, whereas full populations include both monomeric and associated states. Mean profiles and SEs were estimated from nine 200-ns blocks extracted from the final 600 ns of three independent simulations. For the other analogs, monomeric and full-population PMFs showed no substantial differences within statistical uncertainty.

### Section S3 Ring orientations

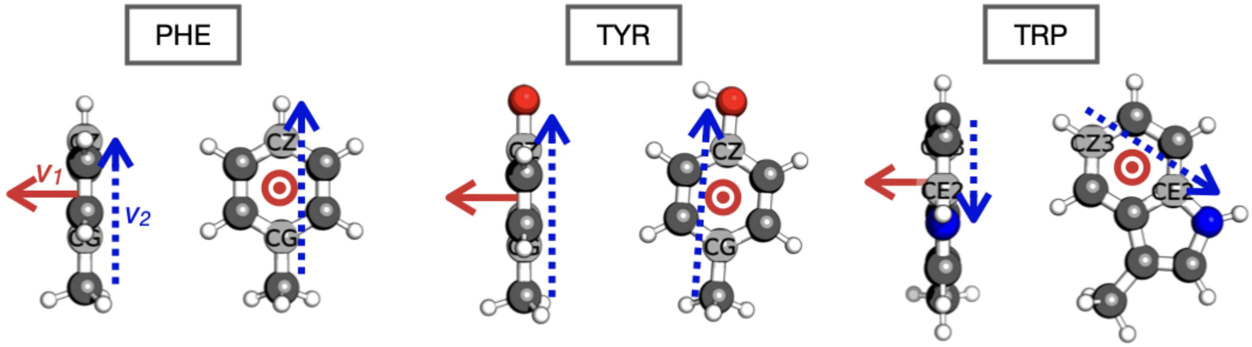

**Figure S8**: Schematic representation of the vectors used to define the ring orientation angles $\theta_1$ and $\theta_2$ for aromatic side-chain analogs. The solid red arrow indicates the vector normal to the aromatic ring plane, and the dotted blue arrow indicates the in-plane reference vector. For PHE and TYR, $\theta_2$ is defined between atoms CG and CZ; for TRP, it is defined between atoms CZ3 and CE2. Molecular structures generated with PyMOL[4] are shown from two perspectives separated by a 90° rotation around the $z$ axis, and vectors pointing out of the page are indicated by $\odot$.

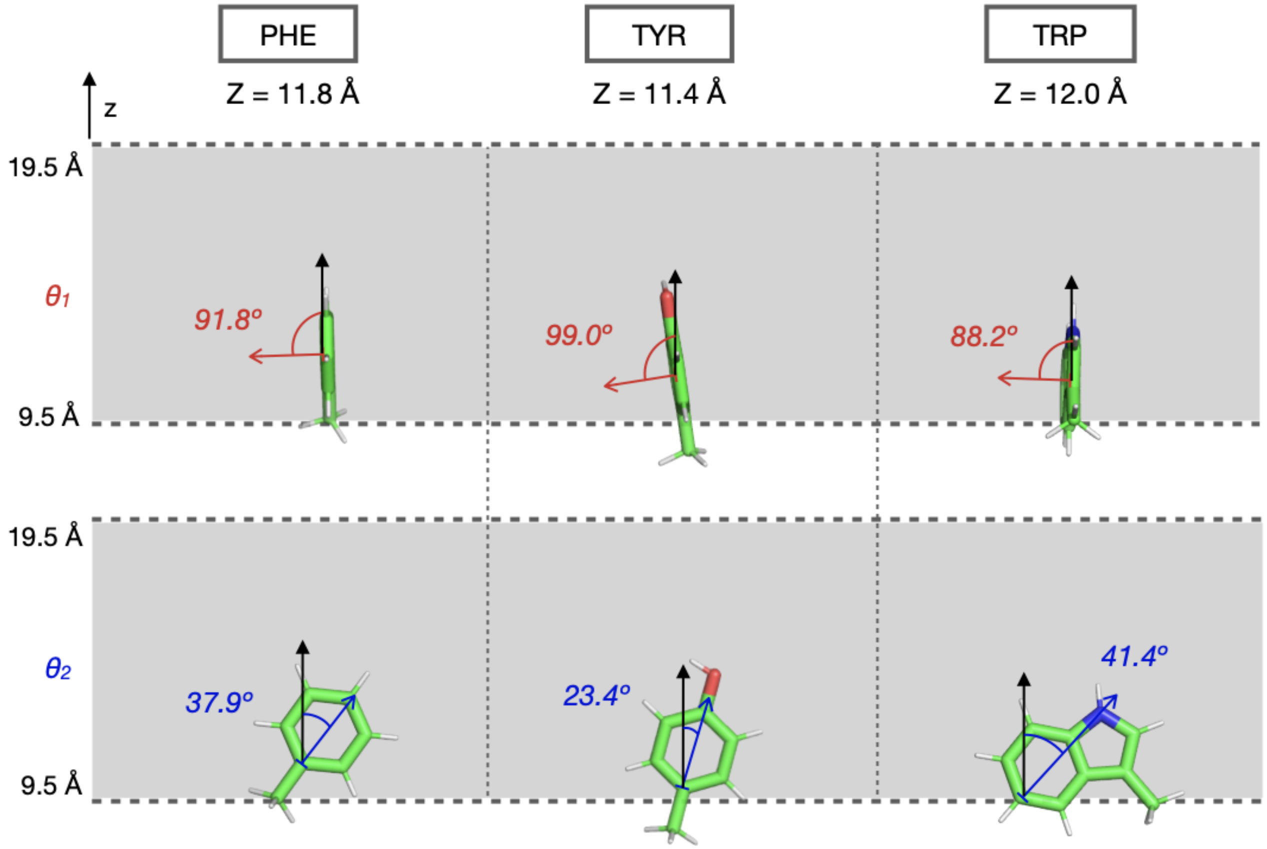

**Figure S9**: Dominant orientations of aromatic side-chain analogs in Region II of the POPC bilayer. Molecular renderings show the most frequently sampled configurations. The gray shaded region between dotted lines indicates Region II, and the arrows indicate the bilayer normal ($z$ axis; black), the ring normal defining $\theta_1$ (red), and the in-plane reference vector defining $\theta_2$ (blue). Molecular structures were generated with PyMOL[4].

### Section S4 Hydrophobicity scales

Computed PMFs were compared with two experimental hydrophobicity scales derived from side-chain partitioning into lipid bilayers. The first is the whole-residue interfacial hydrophobicity scale of Wimley and White (1996), obtained from the partitioning of host-guest pentapeptides (Ace-WL$_X$LL) between water and the POPC bilayer interface at pH 8.0[5]. The second is the side-chain hydrophobicity scale of Moon and Fleming (2011), based on water-to-bilayer transfer free energies measured for OmpLA A210$X$ mutants inserted into the hydrophobic core of a DLPC bilayer at pH 3.8[6].

For comparison with the Wimley-White scale, transfer free energies were extracted from the PMF profiles at the bilayer interface, near the boundary of Regions II and III ($z = 19.5$ Å). For comparison with the Moon–Fleming scale, transfer free energies were taken near the bilayer center, representative of transfer into the hydrophobic core. Figure **S10** compares these computed values with the corresponding experimental data (panel A: Moon–Fleming core scale; panel B: Wimley–White interfacial scale).

Clear correlations are observed between experimental transfer free energies and the PMF-derived values (Pearson correlation coefficient $R = 0.87$ for Moon–Fleming and $R = 0.85$ for Wimley–White), indicating a consistent relationship between computed and experimental partitioning trends. Despite differences between model systems, namely isolated side-chain analogs in the present work and residues embedded in peptides or membrane proteins in the experimental datasets, the level of correspondence is similar for the two scales.

In the Moon–Fleming dataset (Fig. **S10**, panel A), ASP and GLU were considered in their protonated states to reflect the experimental pH (3.8). The data separate into two main groups: aliphatic residues with favorable transfer free energies and polar or charged residues with unfavorable transfer free energies. Given the sampling strategy used in this work, polar residues exhibit larger statistical uncertainty at the bilayer center, which may contribute to their larger deviations relative to aliphatic residues.

In the comparison with the Wimley–White interfacial scale (Fig. **S10**, panel B), MET, ARG, and GLU exhibit the largest deviations, with more favorable insertion free energies than predicted by the experimental scale. MET displays a stronger preference for the bilayer center than for the interface, consistent with its hydrophobic character. For the charged residues ARG and GLU, the more favorable computed values may reflect local bilayer deformations that stabilize the side chains in the isolated-analog simulations; such deformations are more constrained when the residue is part of a short peptide, as in the Wimley–White experiment.

In both experimental benchmarks, side chains are influenced by their local peptide or protein environment. Such context-dependent stabilization or destabilization may shift apparent transfer free energies relative to those of isolated analogs.

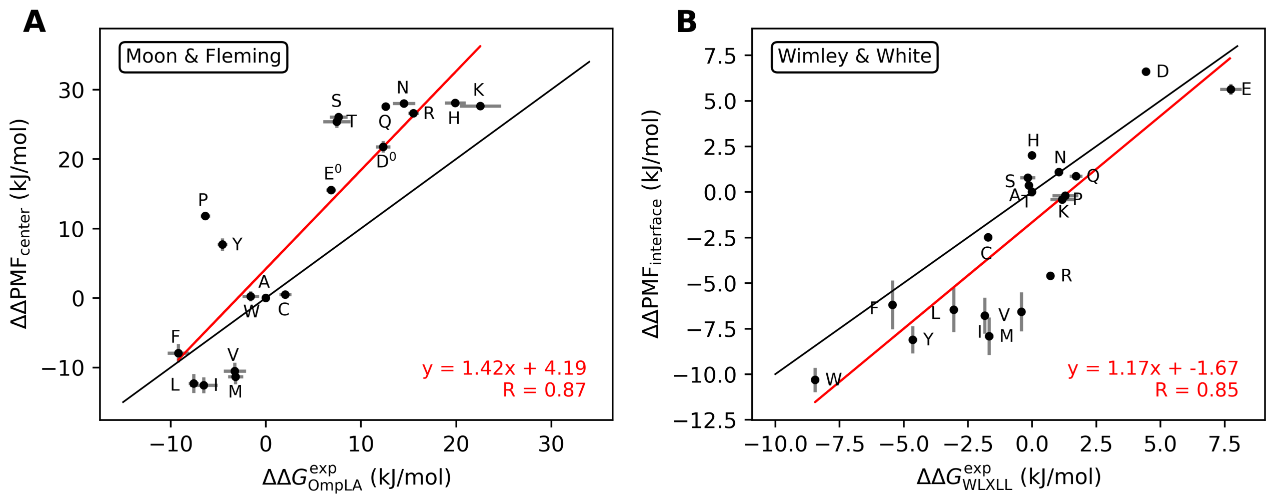

**Figure S10**: Comparison of computed water-to-bilayer transfer free energies for amino acid side-chain analogs with experimental hydrophobicity scales. Symbols show computed versus experimental values for each residue, with vertical and horizontal error bars indicating simulation and experimental uncertainties, respectively. Red lines show linear regressions, and the solid black line indicates perfect agreement ($y = x$). Computed values were extracted from PMF profiles near the bilayer center ($z = 0.5$ Å) in panel A and at the bilayer interface ($z = 19.5$ Å) in panel B. Vertical SEs were estimated from nine 200-ns blocks extracted from three independent simulations. Panel A compares with the Moon–Fleming side-chain hydrophobicity scale, and panel B with the Wimley–White whole-residue interfacial hydrophobicity scale. Residues are labeled with one-letter codes. ASP and GLU were treated in their protonated forms for the Moon–Fleming dataset.
